# Designing a Study of Correlates of Risk for Ebola Vaccination

**DOI:** 10.1101/397547

**Authors:** M. Elizabeth Halloran, Ira M. Longini, Peter B. Gilbert

## Abstract

The rVSV Ebola vaccine was shown to be very efficacious in a novel ring vaccination trial in Guinea. However, no correlates of vaccine protection have been established for Ebola vaccines. Several Ebola vaccine candidates are available, but conducting randomized trials of additional candidates in outbreak situations has become difficult. Establishing correlates of vaccine protection would be useful in helping vaccine candidates become licensed. In this note, we explore power and sample calculations to study potential correlates of risk (protection) during an Ebola vaccination campaign in an outbreak situation under a number of assumptions. At an overall vaccine efficacy of 75%, 50 Ebola endpoints in the vaccinees provided good power. At an overall vaccine efficacy of 90%, 20 Ebola endpoints gave good power under certain assumptions. In the May – July 2018 Ebola outbreak in DRC, over 3000 individuals were vaccinated, with no reported cases in vaccinated individuals. To be feasible, this type of study need Ebola endpoints in vaccinated individuals.

## 1 Introduction

The Ebola Virus Disease (EVD) outbreak in the Democratic Republic of Congo (DRC) of May-July 2018 was the first large outbreak since the EVD outbreak in West Africa in 2014. As of July 24 when Ministry of Health of the DRC declared the outbreak over, there had been 54 cases (38 laboratory confirmed) and 33 deaths reported (External sit rep 17, July 25, 2018). Ring vaccination using the Merck rVSV-Ebola vaccine was implemented in May. In ring vaccination, the contacts and the contacts of contacts (C & CC) of confirmed cases are vaccinated. Also health care workers (HCW) and front-line workers (FLW), such as people working on safe burials were vaccinated. Over 3300 people were vaccinated in that outbreak, with no Ebola cases reported in vaccinated individuals. An Ebola outbreak in and around the North Kivu Province of DRC was reported in the last days of July. As of August 21, 2018, 102 cases (75 confirmed) and 59 deaths (32 confirmed) had been reported (August 21, 2018, Press release, Ministry of Health, DRC).

The rVSV Ebola vaccine was shown to be very efficacious in a novel ring vaccination trial in Guinea [1, 2]. During the vaccine trial in Guinea, not a single case occurred in the arm randomized to immediate vaccination more than 10 days after vaccination. The point estimate of vaccine efficacy was 100% with a lower 95% confidence bound of 70%. Although the vaccine may not be 100% efficacious, it is likely highly efficacious.

Several other Ebola vaccine candidates are available [3]. However, because one vaccine has already been shown to be highly efficacious, it is difficult in an outbreak situation to use a different vaccine that has not been shown in Phase III trials to be effective. It is possible to randomize the different vaccines in non-outbreak situations. Such a trial is being conducted in Guinea, Liberia, Sierra Leone and Mali (Partnership for Research on Ebola VACcinations (PREVAC), ClinicalTrials.gov Identifier: NCT02876328). Although, because there is no Ebola transmission, there have been no Ebola disease events in that trial so far, much new immunogenicity and safety data are being gathered. To guide vaccine candidates through licensure, it would be helpful to establish the immune correlates of risk and protection. Aside from possibly using animal challenge models, such a study must occur in an outbreak situation where vaccine breakthrough events could happen.

In this note, we outline a study of potential correlates of risk (protection) during an Ebola vaccination campaign in an outbreak situation. If the number of cases accrued in the vaccinated individuals is not high enough to reach statistical significance in one outbreak, then the study could be conducted over multiple outbreaks. Ebola is now being viewed as a disease that will return again and again, so that we take a long-term perspective on planning to accrue the information needed to aid licensure of Ebola vaccines.

Here we apply previously developed methods for power calculations [4] for assessing correlates of risk from vaccine efficacy trials or prospective cohort studies to this Ebola vaccine setting. This method to establish correlates of protection relies on the vaccine efficacy varying as a function of the potential immune correlate(s), with vaccine efficacy being higher in those individuals with a measured marker value indicating a stronger immune response and vaccine efficacy being lower in those with a measured marker value indicating a weaker immune response. Under a number of assumptions, the methods can be applied in outbreak situations with no placebo group in contrast to methods developed for traditional prospective placebo controlled trials. We consider power and sample size calculations and discuss the feasibility of conducting such a study.

## 2 Methods

The methods described here are based on the power and sample size calculations for assessing correlates of risk in clinical efficacy trials developed by Gilbert et al [4]. A correlate of risk (CoR) is an immune response that correlates with the clinical endpoint within a cohort such as the per-protocol vaccinated group in a vaccine efficacy trial [5]. A correlate of vaccine efficacy (VE) is an immune response biomarker in vaccine recipients that correlates with VE [6], where VE measures reduction in disease risk in a vaccinated group compared to a comparable placebo or control group.

The approach is based on a nested case-control sampling plan for assessing a CoR in vaccine recipients. It is assumed that a blood sample is drawn, ideally from everyone, at some pre-specified time after vaccination. These blood samples are then stored. It could be that several blood draws are planned, including before vaccination at baseline time 0 (Visit 0). At a minimum, one blood draw at a pre-specified interval after the last vaccine dose, say Visit 1, is drawn and stored. A ‘Visit 1 marker’ may be defined using any data collected up to Visit 1. So, it could be referring to a fold-rise titer since baseline (Visit 0), a very commonly studied marker. Then after the period of follow-up, or at the end of the outbreak, the immune assay(s) will be done on all cases that occurred after the pre-specified marker measurement. Controls will be sampled from those individuals who complete follow-up free of the Ebola endpoint. Simulations have shown that a ratio of controls to cases of 5:1 is efficient [7].

The statistical analysis estimates relative risk (RR) of the Ebola endpoint occurring after Visit 1 through to the end of the follow-up time, contrasting vaccine recipients with different values of the immune response marker. The analysis can be done assuming a continuous marker or a trichotomous marker. The *overall VE* is defined to be the average VE among all vaccinees, whereby some subgroups may have lower or higher VE than the overall VE. Henceforth we use ‘marker’ to mean an immune response marker defined by Visit 1.

Factors that influence the ability to discover and characterize CoRs:

- Number of Ebola endpoint cases between Visit 1 and the specified follow-up period with Visit 1 immune response markers measured.
- Level of overall (average) vaccine efficacy (VE) post Visit 1.
- Strength of the association of the Visit 1 marker in vaccine recipients with the Ebola endpoint and with VE
- Biologically relevant dynamic range/ inter-vaccinee variability of the Visit 1 marker(s).
- Precision of the assay for measuring the functionally relevant immune response (*ρ*).

**–** Noise sources include technical assay measurement error and inter-vaccinee variability in the timing of sampling of immune response markers relative to the immunizations.

The parameter *ρ* is the fraction of the variability of the measured biomarker that is potentially biologically relevant for protection, and is specified to reflect the quality of the biomarker. It is a function of the true variability of the biomarker and variability due to measurement error. A *ρ* equal to 1 indicates there is no measurement error, just true error. If *ρ <* 0.5, likely the biomarker would be a poor correlate.

### 2.1 Continuous marker

For a continuous marker, we are interested in estimating the relative risk as it varies over values of the marker, denoted by *x*:

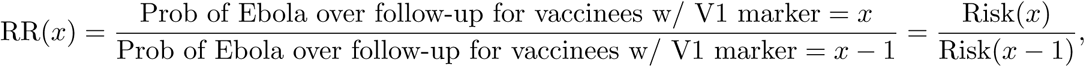

where V1 is Visit 1. A common approach to estimate the relative risk for a rare event setting is to use a logistic regression model designed to handle the nested case-control sampling of the markers [8]. It is also important to assess Visit 1 individual-level signatures, that is, to build models that give the best individual-level classification of whether a vaccine recipient experiences the Ebola endpoint, called individual-level signatures of risk. This can be accomplished using machine learning [9]. We assess a CoR in vaccinees by testing *H*_0_: Risk(x) = Risk for all *x* vs. Risk(x) varies in *x*. We use a 1-sided 0.025-level Wald test [8].

### 2.2 Correlates of VE

CoRs and individual-level signatures in vaccinees are not of direct interest in themselves. We really seek correlates of VE and/or partially valid surrogate endpoints of Ebola disease. Correlates of VE and surrogate endpoints cannot be directly assessed without a placebo/control group. What value is there in CoR/signature assessment in a vaccine group alone?

Identified strong CoRs and signatures in a vaccine group are still very useful, because they may be reasonably hypothesized to also be correlates of VE and/or partially valid surrogate endpoints. Here we illustrate how a CoR in vaccinees translates to a correlate of VE. A correlate of VE is a Visit 1 marker with variability of VE(x) in x defined as

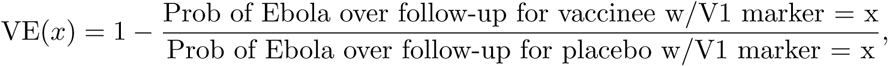

where the V1 marker is the immune response value if assigned to the vaccine group.

The analysis of CoRs/signatures studies the numerator of VE(x) with a strong CoR defined by the numerator strongly varying in *x*. The following assumption implies that a CoR is also a correlate of VE:

- After controlling for the baseline prognostic factors of Ebola disease that are included in the analysis, the placebo-group Ebola risk (denominator) does not vary in *x* as strongly as the vaccine-group risk (numerator).

The stronger the CoR in vaccinees, the greater the credibility of this assumption. Here we present power calculations for a CoR under this assumption that allows it to be interpreted as a correlate of VE.

We can, at some specified month, study a normally distributed quantitative marker with a lower limit of quantification (LLOQ) subgroup. For example, for an overall VE=75%, the lowest 20th percentile of responses might be assumed to have VE=0%. Calculations assume there is a hypothetical placebo group that would have a specified risk of the Ebola endpoint. Calculations are done under the simplifying assumption that VE(x) = 1 − Risk(x Vaccine)/Risk(Hypoth. Placebo).

### 2.3 Trichotomous efficacy and biomarkers

This approach assumes that the vaccinated can be divided into three groups by whether they have low, medium, or high categorical immune response marker(s). It also assumes that the vaccinated can be divided into three groups by their level of vaccination efficacy, whereby they could all be equal, which would be bad for the power.

For a trichotomous biomarker, the correlate of risk (CoR) effect size is defined as the relative risk of disease in the high response group compared to the risk of disease in the low response group. A value of 1 for the CoR effect size means there is no difference in the risk in the two groups, so there is no power. A smaller value of the CoR effect size gives higher power.

For estimation in the trichotomous scenario, the recommended approach is to estimate each probability (numerator for High marker; denominator for Low marker) separately, where for each, one estimates the probability that the failure time is less than some fixed end of follow-up time, accounting for the right-censoring and nested case-control sampling, and adjusting for baseline prognostic covariates by averaging over the covariate-conditional risks. One could also use logistic regression without the averaging, for simplicity. Note that covariate adjustment under logistic regression is not getting at the same estimand as the approach that averages over the covariate-conditional risks.

There are constraints on what sets of parameters are allowed in the power calculations for a trichotomous immune marker. For the trichotomous marker power calculations the user specifies five input parameters relevant for the constraint, where these five parameters determine two other parameters mathematically. The five user-specified parameters are: 1) Overall VE for disease events after Visit 1; 2) A range of VE low response subgroup values; 3) A range of VE medium response subgroup values; 4) The fraction of vaccine recipients in the low response subgroup; 5) The fraction of vaccine recipients in the high response subgroup. From these, the fraction of vaccine recipients in the medium response group, and the range of VE high values, are mathematically determined. One constraint is that all of the VE high response subgroup values must be bounded above by 1.0, (that is, the relative risk cannot be negative), but depending on the input parameter choices this may not occur - in which case one changes the inputs to ensure the VE high values are always *≤*1.

In this paper, the vaccine efficacy in the middle VE group is assumed to equal the overall VE.

To interpret the CoR as a correlate of VE, the risk in the unvaccinated is assumed to be the same for all unvaccinated. Calculations are done under the following simplifying assumptions:

- VE(Low) = 1 − Risk(Low Vaccine)/Risk(Hypoth. Placebo)
- VE(Medium) = 1 − Risk(Medium Vaccine)/Risk(Hypoth. Placebo)
- VE(High) = 1 − Risk(High Vaccine)/Risk(Hypoth. Placebo)

We assess a CoR in the vaccine arm by testing *H*_0_: Risk(Low) = Risk(High) vs. *H*_1_: Risk(Low) */*= Risk(High). We use a 1-sided 0.025-level Wald test from the two-phase logistic regression model accounting for the sampling design [8].

### 2.4 Power Calculations

In this section, we present power calculations for CoR analysis in vaccine recipients, with interpretation of the results in terms of correlates of VE under certain assumptions as in Gilbert et al [4]. We assume a total sample size of *N* = 500, 1000, or 2500 vaccine recipients with blood storage at baseline and Visit 1. Visit 1 would be at whatever time is determined to be a good time to measure the marker(s). This should be standardized across outbreaks. The cases are those who develop Ebola disease after Visit 1 during the follow-up period. The controls are those who complete the follow-up period without experiencing Ebola. After observing the cases and controls, immunological assays would be conducted on the selected (baseline and Visit 1) case-control samples.

We include all cases, that is, vaccinees with the Ebola endpoint, in the analysis. We randomly sample controls, defined as vaccinees free of the Ebola endpoint at the end of follow-up, drawing a 5:1 control:case ratio. Then we calculate power for a Visit 1 CoR for Ebola endpoints after Visit 1.

The power calculations are interpreted assuming that VE is relative to a hypothetical placebo group. For example, VE high = 85% means that vaccinees with a “high” immune response have a 85% reduced rate of the Ebola endpoint during follow-up compared to if they had not been vaccinated. A VE low = 10% means that vaccinees with a “low” immune response have a 10% reduced rate of the Ebola endpoint during follow-up compared to if they had not been vaccinated.

All calculations are done assuming that VE is not below zero (i.e. harmful) for any vaccine recipient defined by *x* continuous or trichotomous. There would be nothing wrong with doing calculations setting VE(Low) = −50%, say, but we are choosing a scenario to study wherein the vaccine does not harm any marker-defined subgroups.

Table 1 summarizes the projected data available for the correlates of risk analysis of vaccine recipients in these power calculations. This assumes an 8% Ebola attack rate in the hypothetical placebo group, and 2% or 0.8% attack rate in the vaccine group (corresponding to 75% or 90% overall VE). At an assumed overall VE = 75%, we present power calculations assuming blood was stored on *N* = 2500, 1000, and 500 vaccinees. The power calculations for overall VE = 90% did not converge when the number of vaccinees with blood stored was assumed to be *N* = 1000 with only 8 expected cases, thus results for overall VE = 90% are presented only for *N* = 2500.

**Table 1:** Projected data available for correlates of risk analysis of vaccine recipients. Assumptions: 8% Ebola attack rate in the hypothetical placebo group; 2% or 0.8% attack rate in the vaccine group (corresponding to 75% or 90% overall VE.)

| Number of Vaccine Recipients with baseline, Visit 1 Blood Storage | Assumed Overall VE | Expected Number Ebola Endpoint Cases with Immunological Data | Number Uninfected Vaccinee Controls (5:1 Ratio) | Total Number with Immunological Data |
| --- | --- | --- | --- | --- |
| $N = 500$ | 75% | 10 | 50 | 60 |
| $N = 1000$ | 75% | 20 | 100 | 120 |
| $N = 2500$ | 75% | 50 | 250 | 300 |
| $N = 2500$ | 90% | 20 | 100 | 120 |

Figure 1 shows the effect on power at overall VE = 75% of varying the fractions of vaccinees with Low and High VE and noise level *ρ* = 0.9 with *N* = 2500 vaccinees with blood storage. The power calculations are done using 1000 simulated data sets, thus the lines are not smooth. The x-axis gives the assumed values of VE high, VE medium, and VE low as they are varied together. On the left, VE high = VE medium = VE low = 75% and power is 0%. On the right, the difference in the vaccine efficacies is most extreme, with VE high = 96%, VE medium = 75%, and VE low = 0%. The results illustrate that power is greater when greater fractions of vaccinees are split into the low and high VE bins and when the difference in VE high and VE low is greater. With 70% of vaccinees at VE high, and 20% of vaccinees at VE low, power of 80% is achieved when VE high *>* 0.83 and VE low *<* 0.50. However, with only 40% of vaccinees at VE high and 10% of vaccinees at VE low, VE high needs to be greater than 86% and VE low needs to be less than 38% to achieve 80% power.

**Figure 1:**
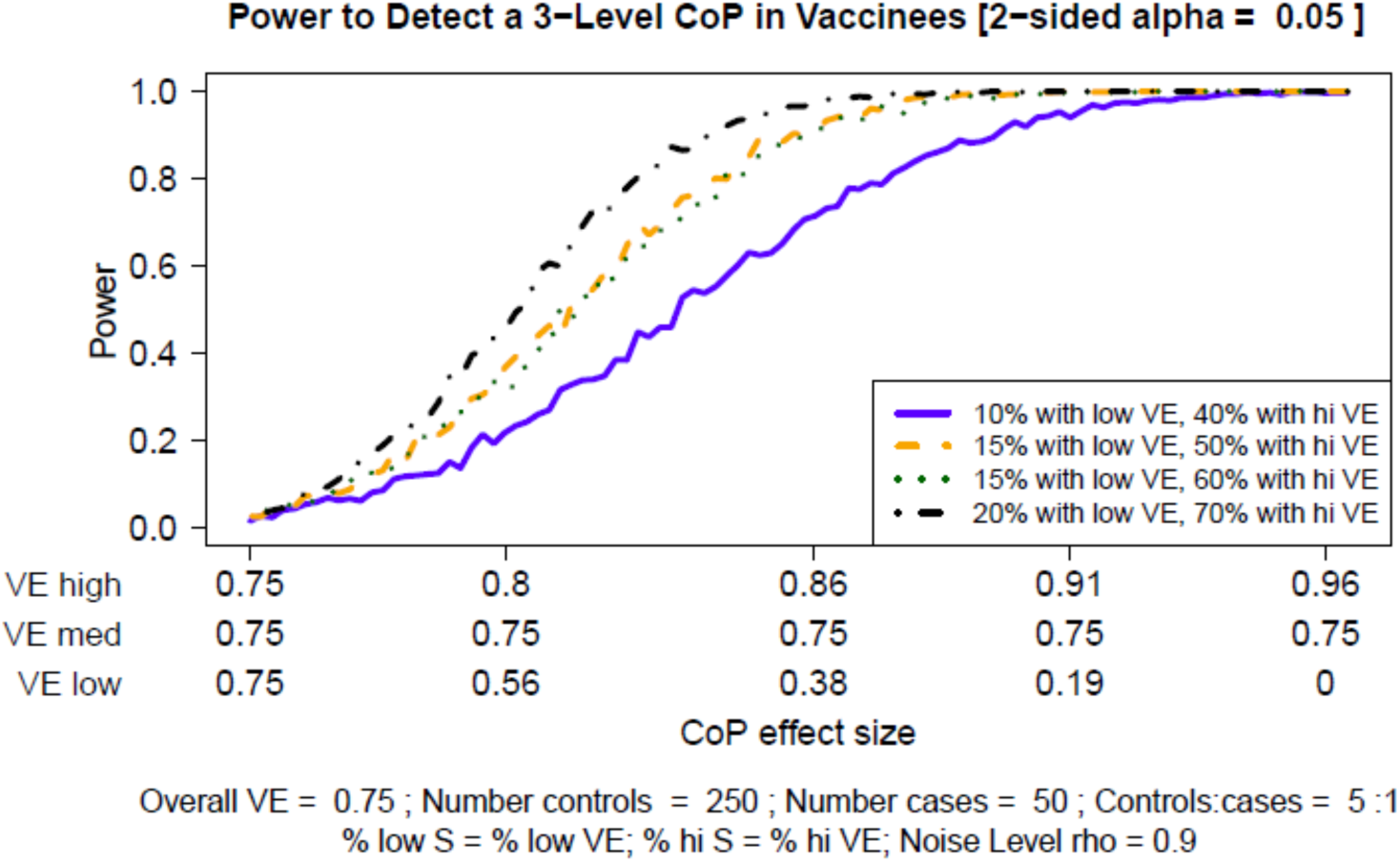
CoR power: Trichotomous marker by fractions of vaccinees with Low and High VE, overall VE = 0.75, at noise level *ρ* = 0.9, *N* = 2500 vaccinees with blood storage.

Figure 2 shows how power varies with the number of vaccinees with blood storage at overall VE = 75% when 70% of the vaccinees are in the VE high group and 20% of the vaccinees are in the VE low group, the most powerful split in Figure 1. Power falls off quickly as the number of vaccinees with blood storage drops. With just 500 vaccinees with blood storage, the difference in VE high and VE low needs to be extreme to achieve 80% power.

**Figure 2:**
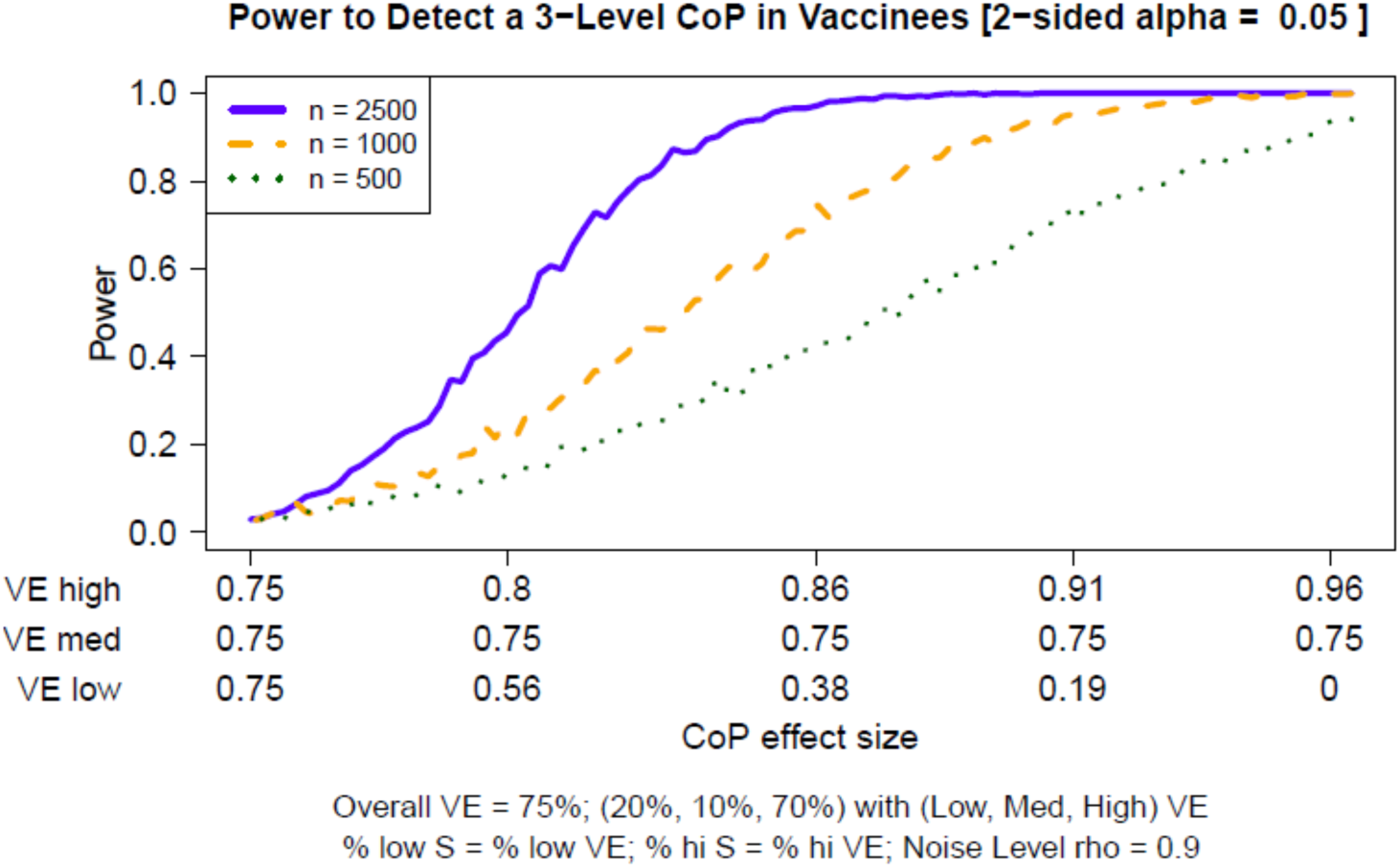
CoR power for a qualifying trichotomous marker at noise level rho = 0.9. Power by numbers of vaccinees with blood storage, overall VE = 0.75. 20% of vaccine recipients have Low response and have VE low. 70% of vaccine recipients have High response and have VE high.

Figure 3 shows the effect on power at overall VE = 90% of varying the fractions of vaccinees with Low and High VE at noise level *ρ* = 0.9. On the left side of the graph, VE high = VE medium = VE low = 0.90 and power is 0. On the right side the VE high = 0.985 and VE low = 0.3, the most extreme values allowed by the constraints. Similar to at overall VE = 75%, the results illustrate that power is greater when greater fractions of vaccinees are split into the low and high VE bins. However, at this very high overall VE there are stricter constraints on what fractions can be in the VE low group as well as on what the lowest VE can be. Here the fraction in the VE low group varies between 5 and 12%, and the fraction in the VE high group varies between 60 and 80%. The VE in the VE low group only goes down to 30%. Power is greater than 80% if about 70-80% of vaccinees are in the VE high group with VE high *>* 94% and 10-12% of vaccinees are in the low VE group with VE low *<* 60%.

**Figure 3:**
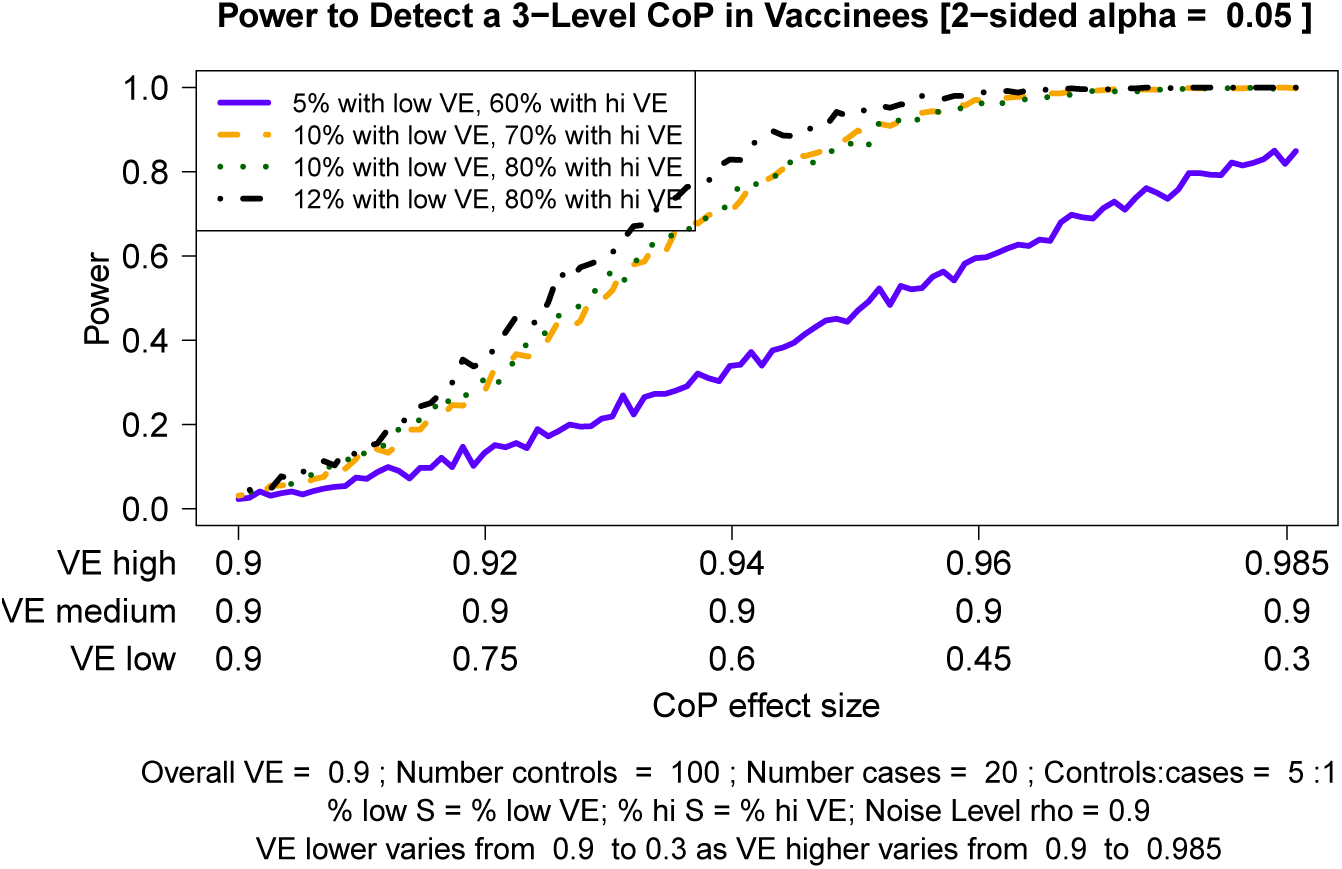
CoR power: Trichotomous marker by fractions of vaccinees with Low and High VE, overall VE = 0.90, at noise level *ρ* = 0.9, *N* = 2500 vaccinees with blood storage.

The figures in the main text all assume a marker with *ρ* = 0.9 marker. This is because (a) additional power calculations provided in the Supplementary Material show that power drops off rapidly with decreasing *ρ*; and (b) in general, it is prudent to allow only immune response markers in a primary correlates study that are known to have a high signal-to-noise ratio, to ensure near maximal power and other reliability properties, and *ρ* = 0.9 is one reasonable benchmark for a qualifying marker.

Additional power calculations are presented in the Supplementary Material. Figures S2 - S8 provide additional power calculations for Overall VE = 75% varying *ρ* and the number of vaccinees with blood stored, as well as presenting results in terms of the CoR relative risk. Figures S9 - S12 present results assuming a normally distributed marker at Overall VE = 75%. Figures S13 - S14 present results varying *ρ* and in terms of the CoR relative risk at Overall VE = 90%.

## 3 Discussion

We have shown that at an overall VE = 75%, 50 Ebola endpoints in the vaccinees provided good power. At an overall VE = 90%, 20 Ebola endpoints gave good power under certain assumptions. In the May – July 2018 Ebola outbreak in DRC, 3300 individuals were vaccinated, with no reported cases in vaccinated individuals. Without events in vaccinated individuals, this type of study is not feasible.

The calculations show that power depends markedly on 1) the strength of association of the Visit 1 marker with the Ebola disease endpoint and that relatively large fractions of vaccine recipients are at the extreme ends of the immune response, 2) the noise level of the marker *ρ*, and 3) the number of vaccine recipients with blood storage. An equally important factor as 1) is

- Factor 1 cannot be controlled; thus we seek to have adequate power for biologically and epidemiologically plausible associations.
- Factors 2 and 3 can be controlled.
- Factor 2: We recommend to include only markers with enough qualification or validation.
- Factor 3: We need to store blood from at least 2500 vaccine recipients to make power adequate for plausible effect sizes. This is especially true when the correlates analysis would consider multiple markers and may need to build in multiplicity correction or training/validation sets.

Factor 1) above can be controlled in a certain sense. If there are animal challenge correlate of protection studies, then one could favor down-selecting immune response markers that showed strong associations with the outcome in the challenge trials. One can use external or previous clinical trials studying immunogenicity of a vaccine candidate, and down-select immune response markers that have the widest biologically relevant dynamic range. Given the numerous immunogenicity studies of current Ebola vaccine candidates, this might be an option. Further, regarding a sampling design point, if there is knowledge on which individuals are expected to have low or high immune response marker values, then one might over-sample individuals based on the predicted extremes to increase power. In practice, however, this might be difficult.

Limitations of our results include that these power calculations all depend on the assumption that the attack rate in the vaccinees would have been 0.08 if no one had been vaccinated, that is, in a hypothetical placebo group. This might be lower or higher. If the attack rate is assumed to be lower or higher, the number of blood samples needed to be stored will increase or decrease about linearly.

Also, the attack rate in the vaccinated could be influenced by indirect effects of ring vaccination. That is, exposure could actually be much lower than if no one had been vaccinated, so the attack rate in the vaccinees may be correspondingly lower, thus a greater number of stored blood would be needed to produce 20 vaccinated cases. The assumption about the hypothetical attack in the absence of vaccination needs to take this into account. Another limitation is that due to the assumed high overall VE (0.75 or 0.90), there are narrow constraints on what proportions can be in the low, middle and high groups in the analysis assuming trichotomous biomarkers.

One complication is that the controls are those who complete the follow-up period without experiencing Ebola. The follow-up period would likely be to the end of the epidemic or possibly when transmission locally has been eliminated. Different outbreaks will have different follow-up periods. The controls for a case from any outbreak would likely need to come from the same outbreak with additional control in the analysis for exposure variables. The consequences of having different follow-up periods in different outbreaks requires further thought.

It is important in the analysis to adjust for variables related to exposure to infection. The reason it is important is that we seek for the correlates of risk to have interpretation in terms of differential biological susceptibility to the outcome, not differential exposure to the outcome. In the ring vaccine trial in Guinea, there were cases in vaccinated people up to six days after they were vaccinated. If early samples from these people had been obtained after vaccination, we would know their biomarker levels as they ramped up around the time they became an early case. This could provide important information about correlates. Going forward, it may be useful to have one or more early samples from individuals.

Having more than one Ebola vaccine available for future outbreaks is essential. Implementing correlates of risk studies for Ebola vaccines should be a public health priority. With a highly efficacious vaccine available, gathering the necessary information will be a challenge.

## Acknowledgments

Funding sources: R37 AI032042 (MEH, IML), U54 GM111274 (MEH, IML), R37 AI054165 (PBG). The content is solely the responsibility of the authors and does not necessarily represent the views of NIH. The code is at http://faculty.washington.edu/peterg/programs.html?

